# Response modality-dependent abstract choice representations for vibrotactile comparisons

**DOI:** 10.1101/802652

**Authors:** Yuan-hao Wu, Lisa A. Velenosi, Felix Blankenburg

**Author notes:** **Corresponding Author:** Yuan-hao Wu, Freie Universität Berlin, Department of Education and Psychology, Neurocomputation and Neuroimaging Unit (NNU), Habelschwerdter Allee 45 14195 Berlin.

## Abstract

Previous electrophysiological studies in monkeys and humans suggest that premotor regions are the primary loci for the encoding of perceptual choices during vibrotactile comparisons. However, these studies employed paradigms wherein choices were inextricably linked with the physical properties of the stimuli and action selection. It raises the question what brain regions represent choices at a more abstract level, independent of the sensorimotor components of the task. To address this question, we used fMRI-MVPA and a variant of the vibrotactile frequency discrimination task which enabled the isolation of choice-related signals from those related to stimulus properties and selection of the manual decision reports. We identified the left, contralateral dorsal premotor cortex (PMd) and intraparietal sulcus (IPS) as carrying information about abstract choices. Notably, our previous work using an oculomotor variant of the task also reported abstract choice representation in intraparietal and premotor regions. However, the informative premotor cluster was centered in the frontal eye fields rather than in the PMd, providing empirical support for a response effector-dependent organization of abstract choice representation in the context of vibrotactile comparisons. Considering our results together with findings from recent studies in animals, we speculate that the premotor region likely serves as a temporary storage site for information necessary for the specification of concrete manual movements, while the IPS might be more directly involved in the computation of choice.

## Introduction

In everyday life, we are continuously encountering situations wherein we need to make decisions based on comparisons between stimuli occurring at different times. Imagine choosing an avocado at a grocery store: one squeezes two or more avocados sequentially and decides for one based on their firmness. The neural processes underlying this type of decision have been extensively studied in the somatosensory domain using the vibrotactile frequency discrimination task (reviewed in Romo & de Lafuente, 2013). In their seminal work, Romo and colleagues trained monkeys to compare frequencies of two sequentially presented vibrotactile stimuli and report with a manual response whether the second frequency (f2) was higher or lower than the first (f1). Crucially, firing rates in premotor regions implicated in the planning and execution of manual movements, such as the supplementary motor area (SMA), ventral (PMv), and dorsal premotor cortices (PMd), have been consistently found to reflect perceptual choices (Hernández et al., 2002, 2010; Romo et al., 2004).

The involvement of motor-related regions during vibrotactile comparisons also agrees well with findings from an influential line of decision-making research in the visual domain. Monkey neurophysiological experiments employing random motion dot tasks with saccade responses consistently reported decision-related signals in regions implicated in saccadic movement (reviewed in Gold & Shadlen, 2007), such as the lateral intraparietal area (LIP, Shadlen & Newsome, 2001; Roitman & Shadlen, 2002), the frontal eye fields (FEF, Kim & Shadlen, 1999; Ding & Gold, 2012), and the superior colliculus (Horwitz & Newsome, 1999; Ratcliff et al., 2003). Findings from these two lines of work have converged to the view that decisions are directly implemented in regions involved in the planning and execution of the resultant action (Gold & Shadlen, 2007; Cisek & Kalaska, 2010). In other words, decisions are implemented in a response modality-dependent manner. Moreover, the posited response modality-specific implementation appears to translate to human vibrotactile comparisons. Herding and colleagues (2016, 2017) reported premotor regions as the most likely source of choice-selective beta oscillatory activity in the EEG signal. The choice-related modulation was localized in the medial part of the premotor cortex when human observers used button presses to indicate their choices (Herding et al., 2016). However, when they reported their choices with saccades, the source of the choice-related modulation shifted to the FEF (Herding et al., 2017).

Of importance, the majority of findings in the context of vibrotactile comparisons were yielded from experimental settings wherein perceptual choices were inextricably linked to the sensory and motor components of the task. In such settings, f1 typically serves as the reference stimulus against which f2 (the comparison stimulus) is compared. Thus, observers will mostly decide for the percept “higher” if frequencies were presented in an increasing order (f1 < f2), and “lower” if presented in a decreasing order (f1 < f2). The abstract contents of perceptual choices are directly bound with the physical properties of the stimulus presentation. Moreover, decisions are typically implemented as choices between two hand or saccade movements so that choosing a particular percept is the same as choosing a specific hand or saccade movement. Due to these dependencies, the presumed choice-related signals may reflect a multiplicity of choice and sensorimotor aspects, rather than the choice per se (Park et al, 2014, see also Huk et al., 2016 for a review). This limitation leaves open the question of whether choices are represented in a more abstract, internal cognitive format, uncontaminated by stimulus order and action selection. For succinctness, we refer to this more abstract type of choice representation as an abstract choice representation throughout the rest of this article.

Our previous work (Wu et al., 2019) addressed this question by means of human fMRI-MVPA and a novel variant of the vibrotactile frequency discrimination task. Intriguingly, although participants’ choices were decoupled from the preceding stimulus orders and ensuing saccade movements used for reporting the decisions, regions implicated in saccade planning and selection such as the FEF and intraparietal sulci (IPS) were identified as representing abstract choices. The finding suggests that activities in these human brain regions are not confined to the sensory and motor aspects of perceptual decisions, but involved in more abstract cognitive computation. Moreover, it hints at the possibility that abstract choices may also be represented in an effector-specific manner.

In the present fMRI study, we sought to further explore the interplay between the topographic organization of abstract choice representations and response modality during vibrotactile comparisons. We asked participants to perform an analogous version of the vibrotactile frequency discrimination task as in our previous work, with saccade responses replaced by manual button presses. Further, the same whole-brain searchlight multivariate analysis routines (Kriegeskorte et al., 2006) as implemented in the previous work was employed to identify brain regions that carry information about abstract choices. Following the interpretation drawn from our previous study, we expected abstract choice representations in premotor regions implicated in the selection of manual responses such as the PMd, PMv, or SMA.

## Materials and methods

### Participants

Thirty-one volunteers participated in the fMRI experiment. They were right-handed, had no history of neurological or psychiatric impairment, and normal or corrected-to-normal vision. Data of four participants were excluded due to poor behavioral performance (accuracy rate < 0.5 in at least one stimulus pair), leaving the data of 27 participants in the analyses (18 females and 9 males; mean age: 25, range: 18–34). All participants provided written informed consent as approved by the ethics committee of the Freie Universität Berlin and received monetary compensation for their time.

### Task design and stimuli

We asked participants to complete a variant of the vibrotactile frequency discrimination task (Fig. 1). Similar to standard versions of the task, participants compared two sequentially presented vibrotactile frequencies and made a decision on whether the frequency of the comparison stimulus was higher or lower than that of the reference stimulus. It differed from standard versions in two important aspects: First, we introduced task rules that alternately designate f1 or f2 as the comparison/reference stimulus across trials so that the perceptual choices were independent of the physical properties of the stimulus order. Second, instead of using a direct choice-motor response mapping, participants reported a match or mismatch between their percept and the proposition of a visual matching cue. After the decision phase, participants selected a color-coded target after a decision phase, from which their perceptual choice was inferred. Hence, participants were not able to plan a specific manual movement or anticipate a target color during the decision phase. As a consequence of these measures, if there were detectable choice-related signals during the decision phase, it would be unlikely to result from the physical properties of the stimulus order or action selection.

**Fig. 1.**
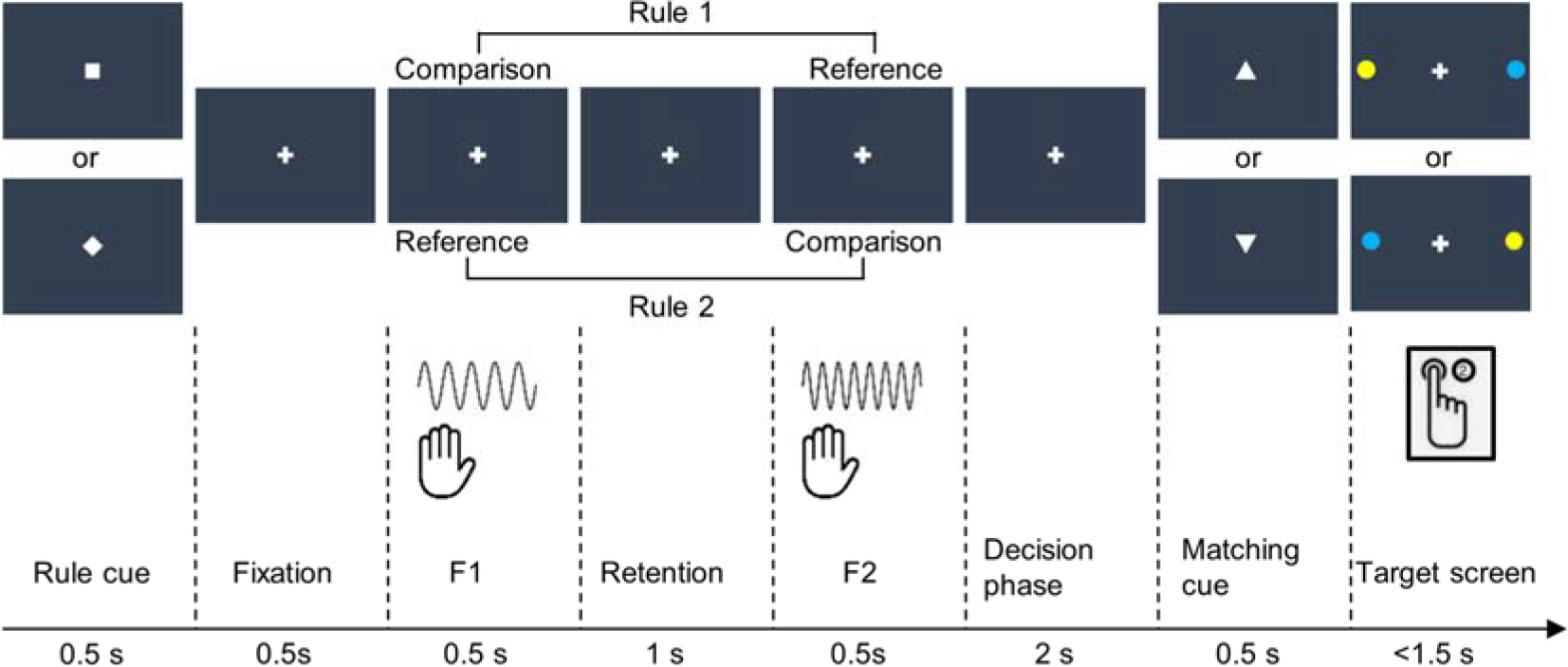
Trial schematic. A rule cue (square or diamond) indicated whether f1 or f2 served as the comparison stimulus. The stimuli presentation was followed by a decision phase. Thereafter, a matching cue (equilateral triangle) was presented. An upward-pointing triangle represented a comparison stimulus of higher frequency, while a downward-pointing triangle represented a lower comparison frequency. Participants compared their perceptual choice with the matching cue. A match or mismatch was indicated by choosing one of the color-coded disks presented in the periphery via a button press. See Wu et al. (2019) for an oculomotor variant of the task.

Each trial was preceded by a variable fixation period (3 – 6 s), during which participants were asked to fixate on a gray cross centrally presented on the screen. The trial started with a switch from the fixation cross to either a square or a diamond for 500 ms, instructing participants which task rule applies in the current trial. In half of the trials, participants used f1 as the comparison stimulus and evaluated whether it was higher or lower than the reference stimulus f2. In the other half, participants made comparisons in the reversed direction. That is, they evaluated f2 relative to f1. The rule cue was followed by two sequentially presented vibrotactile stimuli with different frequencies administered to participants’ left index finger (each of 500 ms separated by a 1 s retention). After a decision phase of 2 s, a visual matching cue in the form of either an upward-pointing or a downward-pointing equilateral triangle appeared centrally on the screen for 500 ms, indicating a comparison stimulus of higher or lower frequency, respectively. Following the offset of the visual matching cue, a target screen with a central fixation cross and two color-coded targets (blue and yellow disks) in the periphery along the horizontal meridian was displayed for 1.5 s. During this time period, participants reported a match or mismatch between their perceptual choice (‘higher’ vs. ‘lower’) and the visual matching cue by selecting one of the color-coded targets corresponding to their report. Depending on the spatial location of the corresponding target, participants pressed the left or right button of a response box held in their right hand with their index or middle finger.

Visual stimuli were generated using MATLAB version 8.2 (The MathWorks, Inc, Natick, MA) and the Psychophysics toolbox version 3 (Brainard, 1997). Except for the two peripheral, color-coded discs on target screens, all other visual symbols were presented centrally in white on a black background. During the fMRI session, visual stimuli were projected with an LCD projector (800 × 600, 60 Hz frame rate) onto a screen on the MR scanner’s bore opening. Participants observed the visual stimuli via a mirror attached to the MR head coil from a distance of 110 ± 2 cm. Suprathreshold vibrotactile stimuli with a consistent peak amplitude were applied to participants’ distal phalanx of the left index finger using a 16-dot piezoelectric Braille-like display (4 × 4 quadratic matrix, 2.5 mm spacing), controlled by a programmable stimulator (QuaeroSys Medical Devices, Schotten, Germany). Frequencies of the first vibratory stimuli (f1) varied between16 and 28 Hz in steps of 4 Hz. The second stimulus was either 4 Hz higher or lower than the preceding f1, yielding a total of eight possible stimulus pairs.

Participants performed six experimental runs of the vibrotactile frequency discrimination task, each lasting ~12.5 min. During each run, each stimulus pair was presented eight times, each time with a unique combination between rule cues (diamond vs. square), matching cues (upward-pointing vs. downward-pointing triangles), and target screens (blue-left, yellow-right vs. yellow-left, blue-right). This yielded a total of 64 trials per run, which were presented in a randomized order. Further, the association between visual symbols and task rules as well as between target colors and match reports was counterbalanced across participants.

### FMRI data acquisition

The fMRI data were obtained with a 3 T Tim Trio MRI scanner (Siemens, Erlangen, Germany) equipped with a 12-channel head coil at the Center for Cognitive Neuroscience Berlin. Functional volumes sensitive to the BOLD signal were acquired using a T2* weighted echo planar imaging sequence (TR = 2000 ms, TE = 30 ms, field of view = 192 mm, flip angle = 70°). Each volume consisted of 37 axial slices and was acquired in an interleaved order (64×64 in-plane, 3 mm isotropic with 0.6 mm gaps between slices). 378 functional volumes were obtained in each experimental run. In addition to the six experimental runs, a T1 weighted structural volume was acquired for co-registration and spatial normalization purposes using a 3D MPRAGE sequence (TR = 1900 ms, TE = 2.52 ms, 256×256 in-plane, 1mm isotropic).

### Data preprocessing and analyses

FMRI data preprocessing and general linear model (GLM) were performed with SPM12 version 6685 (www.fil.ion.ucl.ac.uk/spm) and custom MATLAB scripts (https://github.com/yuanhaowu/DecodingAbstractChoices), while multivariate decoding analyses were performed using The Decoding Toolbox version 3.991 (Hebart et al. 2017, https://sites.google.com/site/tdtdecodingtoolbox/). During the preprocessing, functional volumes were corrected for slice acquisition time differences and spatially realigned to the mean functional volume.

#### Decoding choices

The focus of the present study was to identify brain regions that carry information about choice-related information independent of stimulus order and selection of specific manual response. To this end, we used MVPA combined with a whole-brain searchlight routine to pinpoint brain regions that show distinguishable local activity patterns between different choices during the decision phase.

We first obtained run-wise beta estimates for choice-related activity during the decision phase for each voxel. We fitted a GLM (192 s high-pass filter) to each participant’s data. Separate impulse regressors were defined to model the two choices (‘higher’ vs. ‘lower’), convolved with the canonical hemodynamic function at the onsets of the decision phases. To minimize the number of potential indecisions during decision phases, only correctly answered trials were modelled. Incorrectly answered and missed trials were modelled with a separate regressor of non-interest and not included in the subsequent MVPA. In addition, six movement parameters, the first five principal components explaining variance in the white matter and cerebrospinal fluid signals respectively (Behzadi et al., 2007), and a run constant were added as nuisance regressors, culminating in a total of 120 parameter estimates per participant (20 × 6 runs).

To identify brain regions that exhibit choice-selective activity patterns, a searchlight MVPA was performed on each participant’s data using linear support vector machine classifiers (SVM) in the implementation of LIBSVM 2.86 (Chang & Lin, 2011) with a fixed cost parameter of c = 1. We generated a 4 voxel radius spherical searchlight and moved it voxel-by-voxel through the entire measured volume. The searchlight was centered on each voxel in turn and comprised a maximum of 251 voxels (note that searchlights with 3 and 5 voxel radii yielded similar results). At each voxel, run-wise beta estimates for each of the two choice regressors extracted from voxels within the searchlight formed the 12 response patterns (2 conditions × 6 runs) for the decoding analysis. To avoid overfitting, we estimated the classifier’s decoding accuracy using a leave-one-run-out cross-validation routine. That is, we iteratively trained the classifier to distinguish between response patterns between participant’s choices with data from five runs and tested how well the classifier predicted participant’s choices based on response patterns in the remaining run. This procedure was repeated until all runs were used as the test set. The decoding accuracy of the classifier was estimated as the number of correct predictions divided by the number of all predictions. Decoding accuracy resulting from the searchlight analysis around a given voxel was stored to the corresponding location of a whole-brain volume before the searchlight moved to the next voxel. The searchlight analysis was applied to all voxels in the measured volume so that a continuous whole-brain accuracy map could be obtained. For each voxel in the measured volume, the resulting accuracy map displayed the extent to which the multivariate signal in the local spherical neighborhood was selective to choices. Notably, due to the use of a balanced design, different perceptual choices were expected to have approximately the same number of trials associated with each stimulus order and motor response. That is, each choice regressor contained roughly the same amount of information about stimulus order and button press. Thus, choice-selective activity detected during the decision phase would be unlikely to result from the physical properties of stimulus order or planning of button press responses.

For the group inference, each participant’s accuracy map was transformed to MNI space, resampled to 2 × 2 × 2 mm^3^ voxel size, and smoothed with a 3mm full width at half maximum Gaussian filter. The transformed maps were submitted to a group one-tailed, one-sample t-test to assess whether the decoding accuracy at any voxel was significantly higher than the chance level (50%). A voxel with significant above-chance decoding accuracy would indicate that the local activity pattern around that voxel carries information about choices.

#### Behavioral control analyses

By virtue of the balanced experimental design, the implemented variant of the vibrotactile frequency discrimination task has proven to be capable of disentangling choice-related activity from that related to sensory and motor task components (Wu et al., 2019). However, it remains possible that the classifier could exploit the subtle difference in the distributions of the two stimulus orders (f1 > f2 vs f1 < f1) or motor responses (left vs right button press) between choice conditions to achieve above-chance decoding accuracy (Görgen et al., 2018; Hebart & Baker, 2018). This is of particular relevance for the present study as the balanced number of trials across conditions might not hold after the exclusion of incorrect answered trials and have a biasing effect on MVPA on fMRI data. To address this concern, we applied the same decoding analysis pipeline used with to behavioral data, which enabled us to directly test whether choices can be predicted based on the number of trials associated with different stimulus orders and motor responses in each choice.

For each of the variables of interest, we performed an independent analysis with the following procedure: For each choice in each run, we generated a two-dimensional vector using the number of trials associated with different variable levels. For instance, if a participant responded 15 times with a left and 17 times with a right button press to indicate a comparison stimulus of higher frequency, it was coded as [15 17]. The remainder of the analysis proceeded in a manner analogous to the fMRI data analysis pipeline. Twelve data vectors (2 choices × 6 runs) were used to predict participant’s choices in a decoding analysis with a leave-one-run-out cross-validation routine. For the group inference, we used one-tailed Wilcoxon sign rank tests to probe the statistical significance against chance accuracy (50%). Significant results would imply potential confounds due to the biased distributions of stimulus orders or/and motor responses.

#### Neuroimaging control analysis

As informative clusters identified in the main fMRI analysis include brain regions typically implicated in the planning and execution of manual movements (see result), we did an additional analysis on fMRI data to test whether the result might be confounded with motor planning. We repeated the searchlight choice decoding analysis 100 times for each participant. In each repetition, we randomly sampled a subset of trials so that the number of trials associated with the left and right button presses was fully balanced across choices and runs. We then performed the same GLM and searchlight analysis as described above on a subset of data to obtain a decoding accuracy map per repetition, yielding a total of 100 accuracy maps per participant. The within-participant averaged accuracy maps were then forwarded to a group level t-test to identify brain regions which carry choice-related information. Importantly, by keeping the number of left and right button presses balanced across choices and runs, this analysis eliminated potential confounds related to motor planning. If informative clusters reported in the main result were mainly driven by motor planning rather than by choices, we would not expect choice-related information in the reported regions. Reversely, a similar pattern of informative clusters would strengthen the result of the main analysis.

## Results

### Behavior

The overall behavioral performance of participants during the scanning session was highly accurate. The average accuracy rate was 0.881 (SD: 0.057; range: 0.778 - 0.99), while the average reaction time (latencies between the onsets of the target screens and button presses) was 0.554 (SD: 0.104, range: 0.359 - 0.77).

We further examined participants’ behavioral accuracies and reaction times with three-way repeated measure ANOVAs with task rule (compare f1 against f2 vs f2 against f1), stimulus order (f1 > f2 vs f1 < f2), and f1 magnitude (16Hz, 20Hz, 24Hz, and 28Hz) as within-subject factors, respectively. For the behavioral accuracy, there was no task rule effect observable (F(1,26) = 1.66, p = 0.209). The performance remained stable regardless of which particular rule was applied, suggesting that the cognitive demands were equivalent across rules. In addition, we observed a significant effect of stimulus order (F(1,26) = 7.749, p = 0.001), with a slightly better performance in f1 > f2 trials than in f1 < f2 trials (mean_f1>f2_ = 0.911, mean_f1<f2_ = 0.851, CI_95_ = [0.0166 0.1035]). Moreover, there was a significant interaction between stimulus order and f1 magnitude (F(3, 78) = 11.239, p < 0.001). As indicated by linear trend analyses, participants’ performance decreased slightly with an increasing f1 in f1>f2 trials (slope = −0.0113, p < 0.001), while the performance was unaffected by f1 magnitude in f1 < f2 trials (slope = 0.003, p = 0.233). Contrary to the behavioral accuracy, we did not reveal any difference in reaction times between conditions.

Considering the possibility that response biases and the exclusion of incorrect trials from fMRI analysis may cause differences in stimulus order and motor response distribution between choices and thereby distort the outcome of the fMRI analysis, we performed Pearson chi-square tests on data included in the fMRI analysis, for each participant respectively. The tests did not reveal significant differences in the distribution of stimulus orders and motor responses between choices in any of the participants (all p > 0.1, uncorrected), suggesting that participants’ choice behavior included in the fMRI analysis was not biased by the stimulus order or motor response.

In addition, the same decoding analysis routine as used for the fMRI data was performed to test whether the numbers of trials associated with different stimulus orders and motor responses were predictive of choices. As the results of one-sided, one-tailed Wilcoxon sign rank tests show, neither stimulus order nor motor response was predictive of choices (all p > 0.05, Holm corrected).

Collectively, there is no evidence from our behavioral analyses indicating that the fMRI results reported below were confounded by the physical properties of the stimulus order and selection of the ensuing motor responses.

### Neuroimaging results

The main objective of the present study was to identify brain regions that carry information about perceptual choice independent of the physical properties of stimulus orders and selection of the ensuing manual responses. Using whole-brain searchlight MVPA, we tested for brain regions exhibiting distinguishable local activity patterns between choices during the 2 s decision phase. The result of the whole-brain searchlight analysis is shown in Fig. 3 (displayed at p < 0.05, FDR corrected for multiple comparisons at the cluster level with a cluster-defining voxel-wise threshold of p < 0.001). We were able to decode perceptual choices from the intraparietal sulcus (IPS, mainly in area hIP3; cluster size = 130, peak voxel: [−34 −52 50], t_[26]_ = 5.115, mean decoding accuracy at the peak = 57.737%, CI_95_ = [4.628% 10.847%]) and the dorsal premotor cortex (PMd, BA 6) in the left hemisphere, contralateral hemisphere to the response effector (cluster size = 109, peak voxel: [−20 2 70], t_[26]_ = 4.864, mean decoding accuracy = 60.504%, CI_95_ = [6.066% 14.943%). To test whether choices are indeed represented in a lateralized manner, we conducted two-sided paired t-tests between decoding accuracies extracted from the identified peak voxels and those extracted from the corresponding locations in the right hemisphere (right panel in Fig. 3). These tests show that decoding accuracies extracted from the identified peak voxels were significantly higher than those in the right hemisphere, ipsilateral to the response effector (IPS: t_[26]_ = 2.413, p = 0.002, CI_95_ = [0.928% 11.619%]; PMd: t_[26]_ = 4.43, p < 0.001, CI_95_ = [7.137% 19.467%]), corroborating the lateralized representation of choice-related information.

**Fig. 2.**
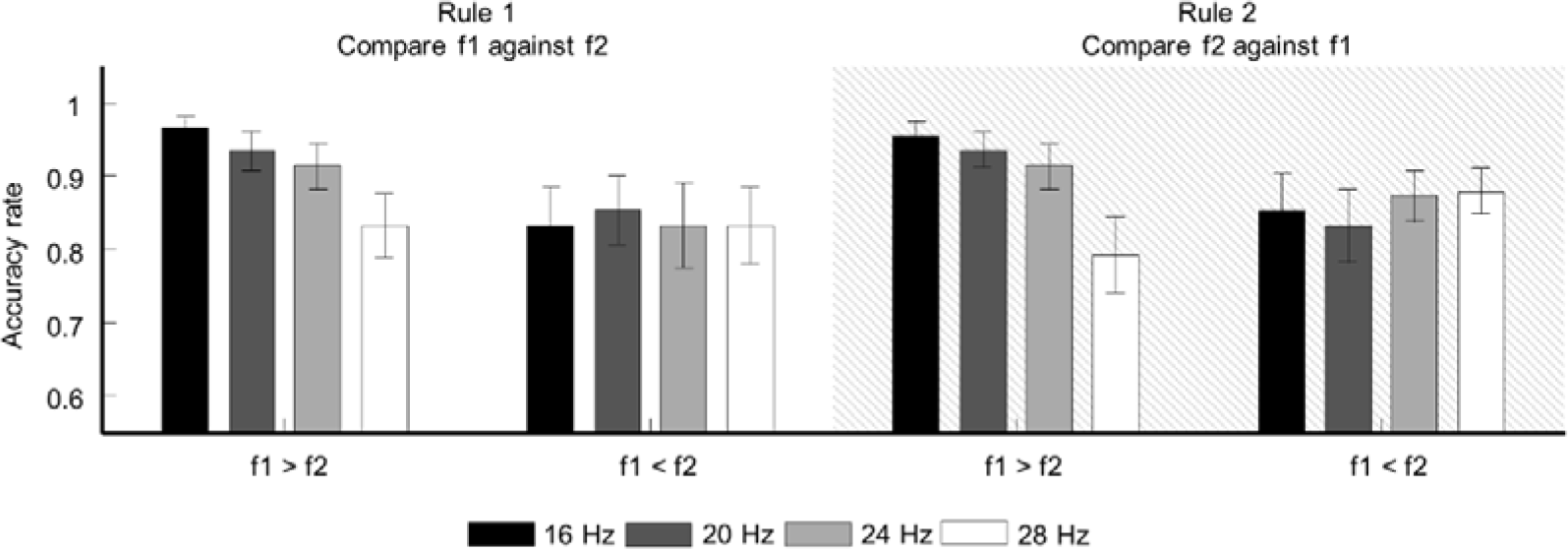
Behavioral performance. The bar plots show the mean accuracy rates across participants over all runs for different stimulus orders, rules, and f1 magnitudes. Error bars represent 95% confidence intervals (CIs) of the means.

**Fig. 3.**
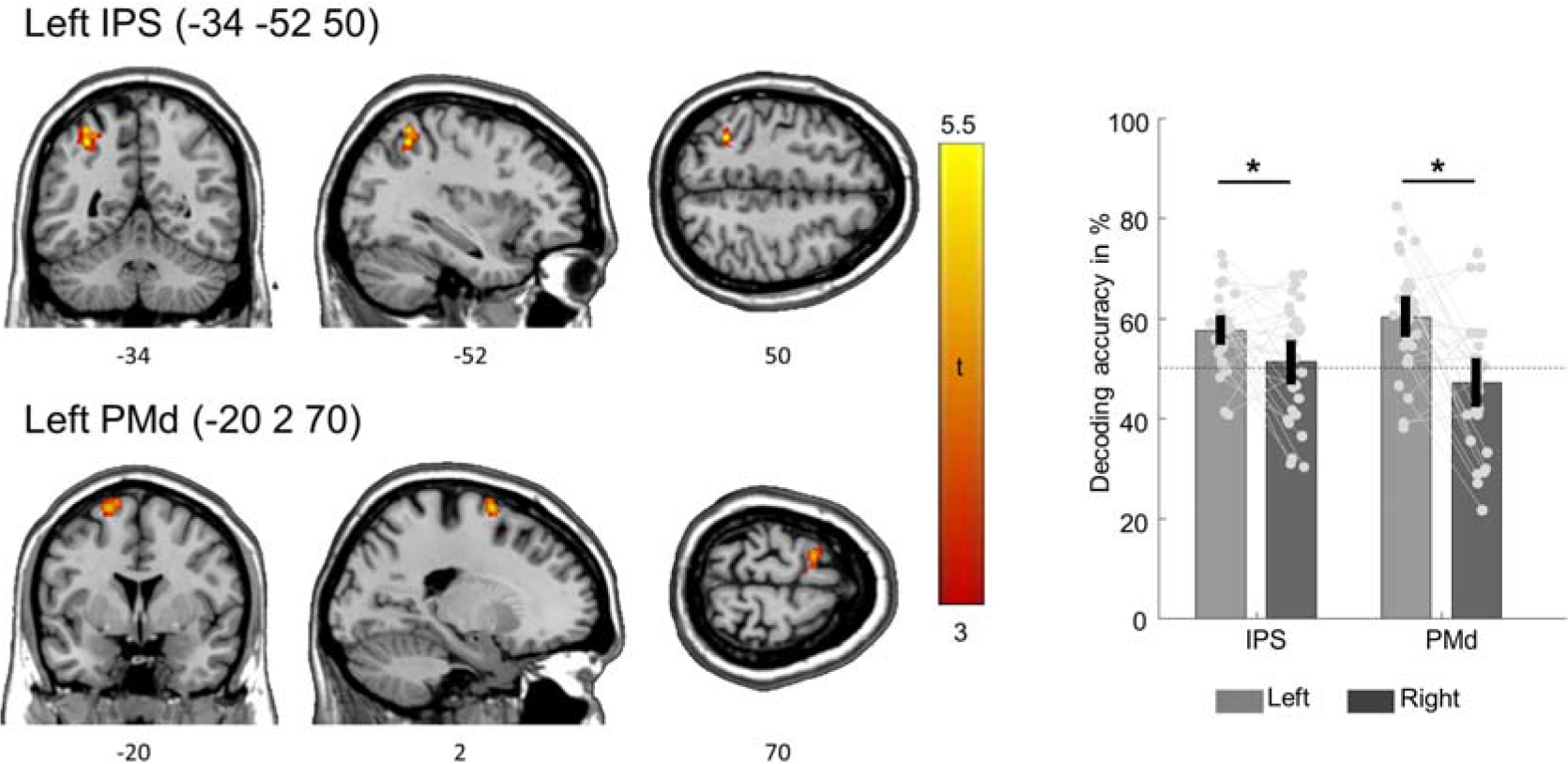
fMRI decoding results. The left IPS and the PMd were found to carry choice-related information independent of stimulus order and ensuing button press, contralateral to the response effector (P_FDR_ < 0.05, cluster corrected for multiple comparisons). Coordinates refer to MNI space and indicate the peak voxel of each region respectively. The unthresholded statistical map can be inspected at https://www.neurovault.org/images/256861/ The bar plot shows decoding accuracies at the reported peak voxels and at the equivalent positions in the right hemisphere, ipsilateral to the response effector. Error bars represent 95% CIs of the means, while dots indicate individual participants’ decoding accuracies in each brain region. Asterisks indicate statistically significant differences between hemispheres at p < 0.05, Holm corrected for multiple comparisons. Participant-specific decoding accuracy maps are available at https://doi.org/10.6084/m9.figshare.9920111.v2

We were further interested in whether decoding accuracies in the reported regions were explanatory to the behavioral performance. To this end, we estimated the Pearson correlation between the decoding accuracy and behavioral performance. We were not able to find statistical evidence for such a linkage between them in any of the reported regions (IPS: rho = 0.089, p = 0.659; PMd: rho = −0.016, p = 0.938).

Importantly, the pattern of informative clusters at the group level remains similar across different searchlight radiuses. We performed the same MVPA with searchlight radii of 3-5 voxels and found that locations of significant informative clusters remain centered in the left IPS and PMd (Fig. 4). Moreover, results of two-sided paired t-tests between all possible pairs show that decoding accuracies do not differ across searchlight radii (all p > 0.05, Holm corrected).

**Fig. 4.**
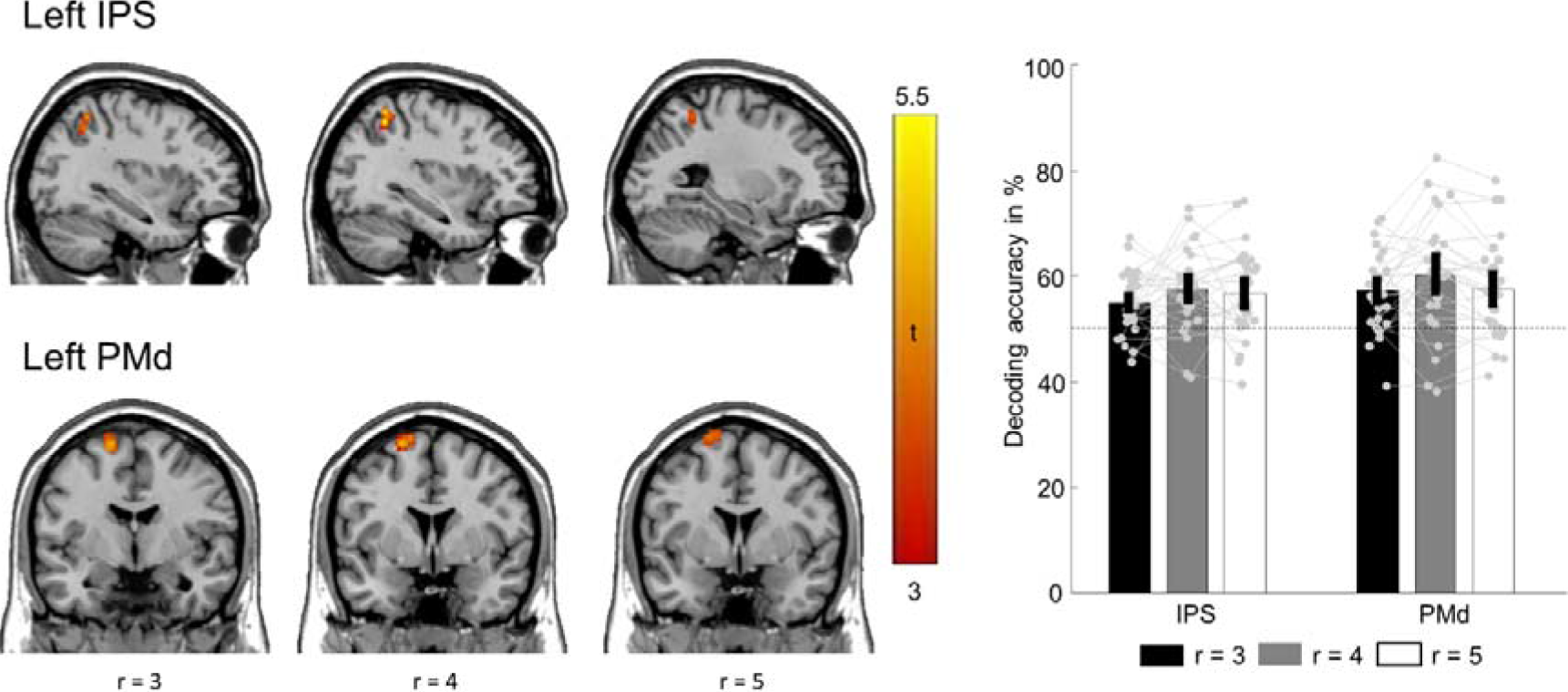
fMRI decoding results using three different searchlight radii. The left panel depicts the informative clusters (one column for each radius, indicated by r). Bar plot in the right panel displays decoding accuracies at peak voxels of the IPS and PMd clusters for each radius respectively. The unthresholded statistical maps are available at https://www.neurovault.org/collections/5936/ Error bars indicate 95% CIs of the means. Grey dots and lines represent individual participants’ decoding accuracies.

We performed an additional decoding analysis to explore whether the identified brain regions with significant above-chance decoding accuracies may result from a bias toward a particular choice-response association. We repeated the searchlight choice decoding analysis and eliminated the potential motor-related confound by keeping the left and right button presses balanced across choices and runs. This analysis yielded a highly similar pattern of brain regions carrying choice-related information as in the main analysis. As shown in Fig. 5 (reported at p < 0.001 uncorrected due to significant reduced amount of data compared to the main analysis), choice-related information was again found in the left IPS ([−34 −52 52], t_[26]_ = 5.173, cluster size = 128, mean = 56.157%, CI_95_ = [3.711% 8.603%) and in the left PMd ([−20 0 72], t_[26]_ = 4.443, cluster size = 76, mean = 57.662%; CI_95_ = [4.117% 11.207%]). Altogether, the results of both behavioral and neuroimaging control analyses suggest that the main results were not driven by motor-related confounds.

**Fig. 5.**
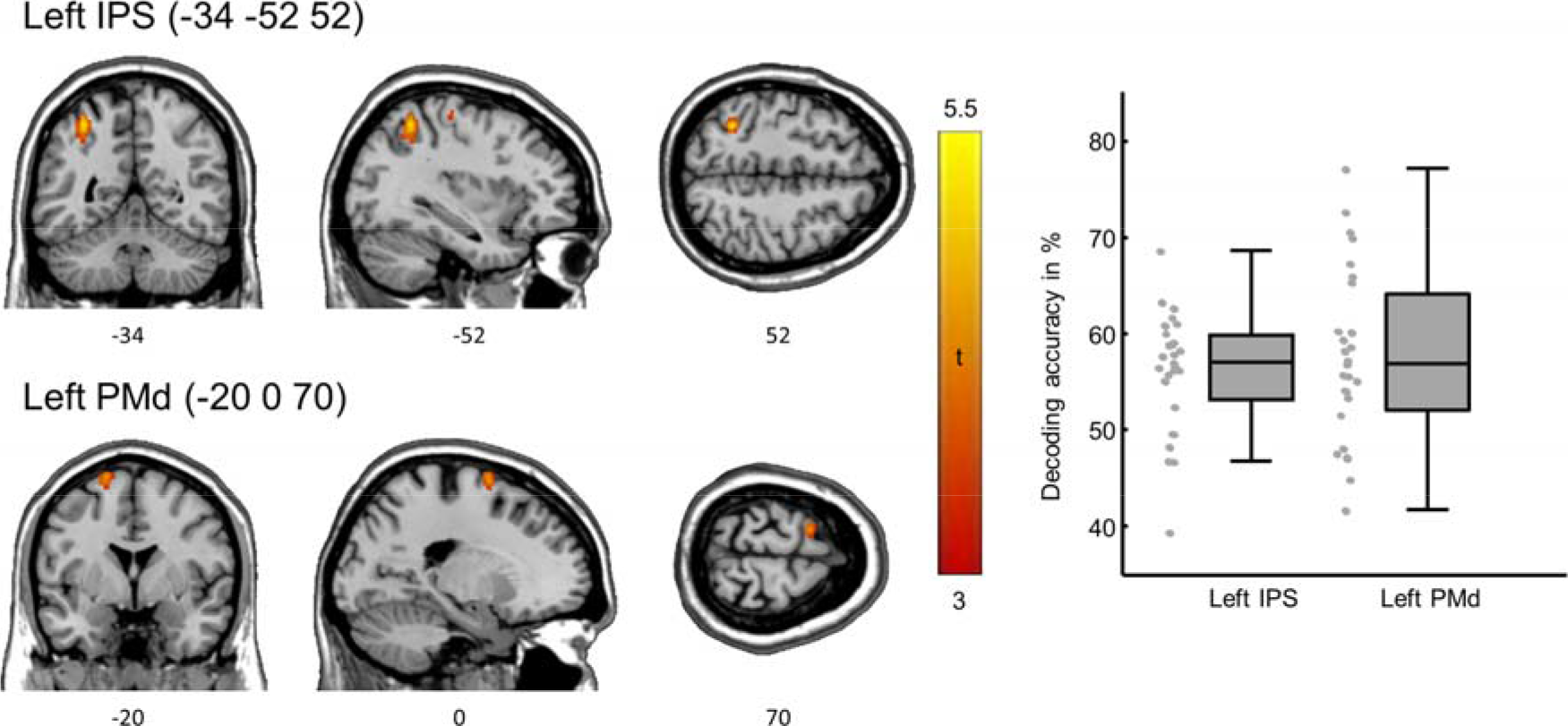
fMRI control analysis result. The left panel displays significant clusters detected by the analysis controlling for motor-related confounds (displayed at p < 0.001, uncorrected). The unthresholded statistical map is available at https://www.neurovault.org/images/256864/ The right panel shows box plots for IPS and PMd separately. Box edges indicate the 25^th^ and 75^th^ percentiles, central horizontal lines correspond to the median. Grey dots represent individual participants’ decoding accuracies.

Next, we compared the result of the present study with that of our previous study, in which decisions were communicated with saccades, instead of button presses (Wu et al., 2019, n = 30). Similar to the present study, choice-selective activity was found in premotor and intraparietal regions, with the difference that it was evident in both hemispheres. The previous study also reported choice-selective activity in the left prefrontal cortex (PFC), while it was absent in the current study. Notably, although both studies identified premotor and intraparietal regions as carrying choice-related information, there were no overlapping clusters. In particular, the premotor clusters identified in the previous study were located in the junction of precentral gyri and the caudal-most part of the superior frontal sulci (peak_left_: [−32 10 62], peak_right_: [34 4 52]), commonly referred to as the FEF (determined with the probabilistic maps by Wang et al., 2015; www.princeton.edu/~napl\vtpm.htm). In contrast, the premotor cluster detected in the current study lies in the adjacent PMd (−20 0 72), dorsocaudal to the FEF (determined with the SPM Anatomy toolbox version 3; Eickhoff et al., 2005), hinting that the location of choice-related information might shift between regions specialized for eye and hand movements depending on what response effector is used.

To further assess this possibility, we ran a set of regions of interest (ROI) analyses. First, we took the peak voxels in the bilateral FEF from the previous study as the ROI for the current data. For each participant, we extracted decoding accuracies from these voxels and averaged them. The averaged decoding accuracies were then submitted to a two-tailed, one-sample t-test against the chance level. Likewise, we used the peak voxel of the PMd cluster from the present study as the ROI for the previous study and tested whether choices could be reliably decoded from the PMd. The results of these ROI analyses support the interpretation of an effector-dependent shift of choice representation within the premotor cortex (Fig. 6). Despite the higher sensitivity of ROI approach, the mean decoding accuracy computed from the bilateral FEF in the present study did not surpass the chance level (t_[26]_ = 1.534, mean = 52.272%; CI_95_ = [−0.772% 5.315%], p = 0.137). Likewise, the mean decoding accuracy in the left PMd derived from the previous study did not differ significantly from the chance level (t_[29]_ = 2.172, mean = 54.301%; CI_95_ = [0.250% 8.352%], p = 0.076, Holm corrected). That is, when manual response was used, choice could only be reliably decoded from the left PMd, but not from the FEF. Conversely, choice could only be read out from the FEF, but not from the PMd, when saccadic response was required (Fig. 6).

**Fig. 6.**
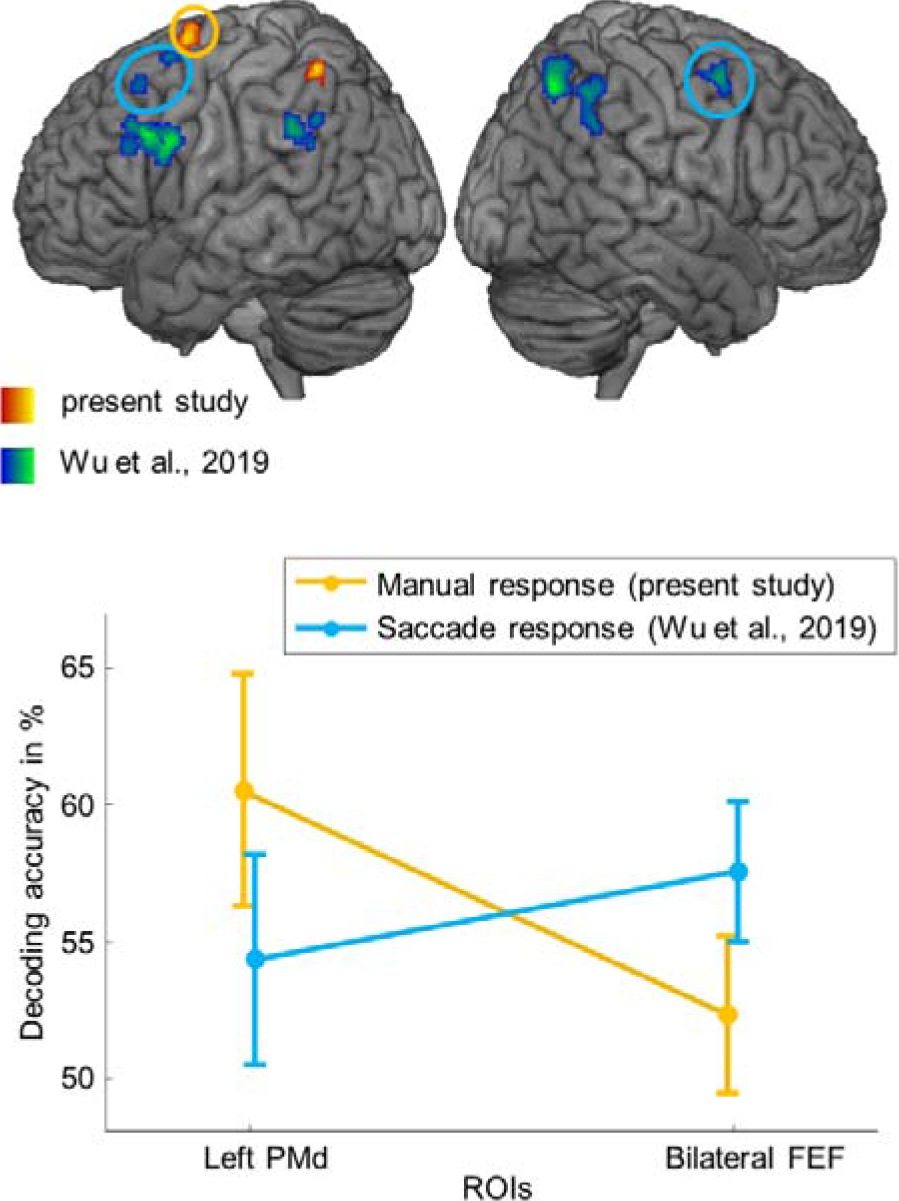
Comparison with results from the saccade version of the task (Wu et al., 2019, n = 30). The upper panel displays brain regions carrying choice-related information as identified in the present study (in red-orange) and those detected in our previous work using saccades as decision reports (in blue-green, unthresholded statistical map available at https://www.neurovault.org/images/63793/), both displayed at p_FDR_ < 0.05, cluster corrected. The circles indicate the premotor and intraparietal clusters used for ROI analysis. The lower panel depicts mean decoding accuracies across participants collapsed across response modalities and effector-specific regions. Error bars indicate 95% CIs of the means.

## Discussion

In the present study, we sought to identify human brain regions that represent abstract choices in the context of vibrotactile frequency comparisons. We used fMRI combined with a variant of the vibrotactile frequency discrimination task which allowed us to dissociate choice-selective BOLD signals from those related to the physical properties of stimulus orders and the selection of manual responses. We identified the left IPS and PMd, contralateral to the response effector, as carrying choice-related information. Notably, using the same task, but saccades as response effector, our previous study (Wu et al., 2019) also reported choice-related information in intraparietal and premotor regions. Interestingly, the informative premotor cluster was centered in the FEF rather than in the PMd. Evidence from these two studies suggests a response modality-specific organization of abstract choice representations in the context of vibrotactile comparisons.

The pivotal role of the premotor cortex in decision formation during vibrotactile comparisons has been established by the seminal work of Romo and colleagues using neurophysiological recordings in monkeys (reviewed in Romo and de Lafuente, 2013). The premotor cortex is strongly implicated in the computation of comparisons between the two sequentially presented stimuli, based on the consistent observation of choice-predictive signals before the initiation of manual responses (Hernández et al., 2002; 2010). In line with these reports, we identified the dorsal part of the premotor cortex as carrying choice-related information, with the crucial difference that choices in the present study were independent of sensorimotor components, while choices in the above-mentioned monkey neurophysiological studies were inextricably linked with them. Taking this into account, the finding of such abstract choice representations in a region that is primarily associated with the planning and preparation of manual actions may not appear straightforward. Indeed, results from few human fMRI studies in the visual domain, wherein perceptual choices were disentangled from specific actions, are inconsistent. On the one hand, several studies failed to find evidence for decision-related BOLD signals in the premotor cortex when choices were decoupled from actions (e.g., Hebart et al., 2012; Filimon et al., 2013). On the other hand, premotor activity reflecting categorical choices regarding the stimulus identity independent of motor planning has been shown in other human fMRI studies (e.g., Hebart et al., 2014). With this study, we provide additional fMRI evidence for a premotor involvement in the representation of choices in a more abstract, internal cognitive format.

Hereof, it is important to note that the analysis we used in the present study does not permit an inference about whether abstract choices are indeed encoded in the PMd or generated elsewhere. Independent of this issue, one possible explanation for our premotor finding is that the PMd serves as a node for short-term storage of abstract choice representations and the transformation into commands for concrete manual movement once all information required for the execution of specific actions are known. This interpretation agrees with a recent study showing a causal role of the premotor cortex in the flexible stimulus-response mapping in mice (Wu Z. et al., 2019) and monkey neurophysiological studies implicating the PMd in the retrieval and integration of task-relevant information necessary for specification of particular actions (e.g., Nakayama et al., 2008; Yamagata, 2009, 2012).

While there is a vast amount of neurophysiological evidence for the premotor involvement during vibrotactile comparisons, neural activities in the posterior parietal cortex (PPC) has remained largely unexplored in this context. Nevertheless, our finding of intraparietal choice representation was not surprising. Similar to the premotor area, posterior parietal regions are thought to be crucially involved in various decision-making tasks, most prominently when decisions are communicated by saccades (Gold & Shadlen, 2007). In particular, activity in the monkey LIP (homologous to the intraparietal subregions in humans) has been shown to mimic the presumed evidence accumulation toward one or the other saccade choices and thereupon regarded as the explicit neural representation of the evolving decisions (Shadlen & Kiani, 2013, but see Huk et al., 2017 for a critical review). Moreover, evidence from recent studies on a wide range of decision-making tasks suggests that PPC’s involvement is not confined to motor decisions but pertains to decisions at different levels of abstraction. For instance, both monkey and human PPC have been shown to represent choices that were independent of the planning of saccade responses (Bennur & Gold, 2011; Hebart et al., 2012). Among studies in the broader context of decision making, findings from monkey neurophysiological recordings using visual categorization tasks are particular revealing (reviewed in Freedman & Assad, 2016). In these studies, monkeys were trained to perform delayed match-to-category tasks in which they decide whether the motion direction of the sample stimulus and the test stimulus belong to the same category based on a previously learned, arbitrarily defined boundary. After the test stimulus, monkeys indicated their decision on a match or mismatch with manual or saccadic responses. Using this task, LIP has been shown to exhibit signals reflecting the categorical choice which cannot be attributed to specific sensory stimulus properties nor action selection (Freedman & Assad, 2006; Swaminathan & Freedman, 2012; Swaminathan et al., 2013). Such categorical information is reminiscent of the choice-related information observed in our study as both are dissociated from sensory and motor components of the task and are thus, represented at a similar level of abstraction. The similarity between them opens the possibility of a common mechanism and thereby boosts the notion of the PPC, and IPS more specifically, as a central node mediating abstract cognitive computations (Freedman & Assad, 2016).

Given the above-mentioned functions ascribed to the PPC, one question which naturally emerges from our results is whether the reported choice-related information is directly computed in the PPC via the evidence accumulation process or other mechanisms. We are not able to answer this question with our experimental design. In this study, we only used stimulus pairs with supra-threshold differences to facilitate the decodability of choice-related information. This is, however, problematic for assessing neural correlates of evidence accumulation as they would, according to the accumulation-to-bound model (Ratcliff et al., 2016), provide strong momentary evidence signals which are difficult to distinguish as such. Similar to the premotor cortex, it is possible that the IPS merely receives choice-related signals from elsewhere in the brain and thus, is not actively involved in the decision formation. However, there is evidence from several lines of research that warrants the IPS being a promising candidate region for decision formation during vibrotactile comparisons.

First, vibrotactile comparisons as implemented in the present study can be regarded as a process in which a choice is made based on the relation between two magnitudes. Combined evidence from monkey neurophysiology and human neuroimaging suggest that magnitudes and the relation between them are encoded by a network comprising the IPS and lateral PFC (reviewed in Jakobs et al., 2013). Moreover, the IPS appears to be the first region within this network to process magnitude information (reviewed in Nieder, 2016). Second, Herding and colleagues (2019) showed that the centro-parietal positivity (CPP) in EEG signal, which has been suggested as a proxy for accumulated evidence across a variety of decision-making tasks (O’Connell et al., 2012; Kelly & O’Connell, 2013), also indexes the amount of sensory evidence during vibrotactile comparisons. More specifically, they identified the left IPS as the likely source of the CPP component reflecting the signed subjectively perceived difference between two frequencies. Notably, in this study, participants always compared f2 against f1. It would be interesting to explore whether and how this effect is modulated by comparisons in the reversed direction. Finally, using a reversible inactivation approach to investigate PPC’s contribution to sensory evaluation and action selection. Zhou and Freedman (2019) revealed that monkeys’ decisions were more severely affected when visual stimuli, rather than motor targets, were placed in the inactivated receptive fields of LIP neurons under investigation, providing compelling evidence for the causal role of the PPC in the sensory aspect of visual decisions. Given that the IPS is thought to have a similar role as a mediating node in the sensorimotor transformation across multiple sensory domains, it is intriguing to see whether a causal effect could also be demonstrated during vibrotactile comparisons.

With the present finding of premotor and intraparietal choice-selectivity, we have also replicated the finding of our previous study using the same task but with saccades as the response modality (Wu et al., 2019). When comparing both studies more closely, two differences are apparent. First, choice-related information was found in bilateral premotor and intraparietal regions when saccades were used. However, when manual responses were required, the premotor and intraparietal selectivity was only evident in the contralateral hemisphere. Moreover, we observed a relocation of choice-related information within the premotor area from the FEF to the PMd. Importantly, we did not assign these functional labels merely based on the required response modalities tasks. Both the FEF and the PMd were determined by means of well-established functional probability maps. In addition, the spatial arrangement of the FEF and the PMd clusters as identified by the spatially unbiased whole-brain searchlight routines in these two studies corresponds well to that reported in monkeys (e.g. Petrides, 1982; Halsband & Passingham, 1982; Bruce & Goldberg, 1985) and humans (Amiez, 2006), with saccade-related premotor region lying more anterior and rostral to premotor region exhibiting activities related to manual movements. Thus, it is unlikely that these differences were merely a by-product of idiosyncratic differences between samples. Altogether, the results from these two studies suggest a response modality-dependent organization of abstract choice representations. One question emerged from this interpretation concerns whether the posited response modality-dependent organization of abstract choice information is confined to a specific level of abstraction. For instance, the dependency observed in our studies might result from the explicit foreknowledge of the required response modality. Evidence from other fMRI studies suggests that decision-related activities may occur elsewhere when the required response modality is not known (Ho et al., 2009; Liu and Pleskac, 2011; Filimon et al., 2013). In this light, future studies combining the present task with a wide range of response modalities, target locations, and task difficulties will provide essential insights into how vibrotactile choices are evolved and transformed into internal cognitive states in humans.

## Acknowledgements

This research was supported by the Deutsche Forschungsgemeinschaft (DFG) – project number 409180874. We thank Pia Schröder for many inspiring discussions and Sam Gijsen for proof reading the article.

## Author contributions

Y.W. and F.B. designed the experiment and interpreted the results. Y.W. and L.A.V. conducted the experiment, Y.W. analyzed the data and wrote the manuscript. L.A.V. and F.B. reviewed and edited the manuscript.

## Competing interests

The authors declare no competing financial interests.

## Notes

https://doi.org/10.6084/m9.figshare.9920111.v2

https://www.neurovault.org/collections/5936/

https://github.com/yuanhaowu/DecodingAbstractChoices

